# Multi-omics profiling reveals phenotypic and functional heterogeneity of neutrophils in COVID-19

**DOI:** 10.1101/2023.09.02.556069

**Authors:** Lin Zhang, Hafumi Nishi, Kengo Kinoshita

**Author notes:** Correspondence: Lin Zhang.

## Abstract

**Background:** Accumulating evidence has revealed unexpected phenotypic heterogeneity and diverse functions of neutrophils in several diseases. Coronavirus disease (COVID-19) can alter the leukocyte phenotype based on disease severity, including neutrophil activation in severe cases. However, the plasticity of neutrophil phenotypes and their relative impact on COVID-19 pathogenesis has not been well addressed. This study aimed to identify and validate the heterogeneity of neutrophils in COVID-19 and evaluate the phenotypic alterations for each subpopulation.

**Methods:** We analyzed public single-cell RNA-seq, bulk RNA-seq, and human plasma proteome data from healthy donors and patients with COVID-19 to investigate neutrophil subpopulations and their response to disease pathogenesis.

**Results:** We identified eight neutrophil subtypes, namely C1–C8, and found that they exhibited distinct features, including activation signatures and multiple enriched pathways. The neutrophil subtype C4 (DEFA1/1B/3+) associated with severe and fatal disease. Bulk RNA-seq and proteome dataset analyses using a cellular deconvolution approach validated the relative abundances of neutrophil subtypes and the expansion of C4 (DEFA1/1B/3+) in severe COVID-19 patients. Cell– cell communication analysis revealed representative ligand-receptor interactions among the identified neutrophil subtypes. Notably, the C4 (DEFA1/1B/3+) fraction showed transmembrane receptor expression of CD45 and CAP1 as well as the secretion of pro-platelet basic protein (PPBP). We further demonstrated the clinical potential of PPBP as a novel diagnostic biomarker for severe COVID-19.

**Conclusion:** Our work has great value in terms of both clinical and public health as it furthers our understanding of the phenotypic and functional heterogeneity of neutrophils and other cell populations in multiple diseases.

## Background

Coronavirus disease (COVID-19), caused by severe acute respiratory syndrome coronavirus 2 (SARS-CoV-2), can alter the leukocyte phenotype in severe cases, including alterations in the immune system, exhaustion of lymphocytes, and changes in hematopoiesis [1–6]. In addition to macrophage, B cell, and T cell activation, numerous studies have shown that neutrophil activation by virus infections [2,7–11]. A recent study identified alterations in neutrophil phenotypes that may be associated with the pathogenesis of several diseases [12]. However, the plasticity of neutrophil phenotypes and their relative impact on COVID-19 pathogenesis have not been well described.

Neutrophil populations are characterized by the variability in their phenotypes and functions. Over the past five years, accumulating evidence has revealed unexpected phenotypic and functional heterogeneity of neutrophils in multiple diseases, including cancers, atherosclerosis, and type-2 diabetes [12]. For example, three subsets of neutrophils, namely mature high-density neutrophils, mature low-density neutrophils (LDNs), and immature LDNs, have been observed in the circulation of both patients with cancer and tumor-bearing mouse models [13–16]. A study of 124 patients and eight mice revealed the heterogeneity of neutrophils in liver cancer, and some of the subpopulations were found to be associated with an unfavorable prognosis, where the CCL4+ neutrophil population can recruit macrophages and the PD-L1+ population displays suppression of T-cell cytotoxicity [17]. Furthermore, neutrophil subpopulations have been identified to have opposing immune functions in some contexts, such as collateral damage during post-injury inflammation, compared with the regeneration of post-injury tissue [18–21]. Several other anti-inflammatory functions, including the suppression of T-cell function and release of α-defensins to inhibit macrophage-driven inflammation, have also been described [12]. Although the role of neutrophils in viral infections remains unclear, increasing evidence has demonstrated that neutrophils contribute to resolving viral infection [22]. In a mouse model, depletion of neutrophils with an anti-Gr-1+ antibody caused severe diseases, such as respiratory dysfunction and dysregulated immunity during influenza A virus (IAV) infection [23,24]. Literature on viral respiratory diseases (VRDs) provides an understanding of neutrophil heterogeneity and functions in several stages, such as heterogeneous neutrophil infection, development, and activation [22]. In the infection stage, IAV-infected neutrophils produce less human cathelicidin LL-37 and can act as antigen-presenting cells for antiviral CD8+ T cells [25,26]. In the development stage, high fractions of immature neutrophils, which was not affected by bacterial coinfection, was observed in infants with various VRDs [27]. In the activation stage, a degranulated form of neutrophils (CD16^Int^ low-density neutrophil population) contributes to immunosuppression in VRDs and acute respiratory distress syndrome [28].

Discoveries of the versatile phenotypes and functions of neutrophils have opened new doors to understand their contributions to antimicrobial functions as well as homeostatic and pathogenic immune processes [29]. The present study aimed to analyze scRNA-seq, bulk RNA-seq, and plasma proteome data of healthy individuals and patients with COVID-19 to identify and validate the heterogeneity of neutrophils. It further investigated the alterations in phenotypes and contributions in response to disease severity for each neutrophil subpopulation. The workflow used for the neutrophil subpopulation investigation and validation is shown in Figure 1. We observed that neutrophils were activated after SARS-CoV-2 infection in the scRNA-seq, bulk RNA-seq, and proteome datasets. First, the standard Seurat workflow was used to identify finer subtypes of neutrophils. Activation signatures of each subtype were detected. Biological functions and cell-cell communication analyses were subsequently used to describe the phenotypic and functional heterogeneity of each subtype. Comparisons between severe COVID-19 and healthy cells were performed to reveal the mechanisms underlying COVID-19 progression, including differentially expressed gene (DEGs) identification and pathway enrichment analysis. Next, the subtypes identified using scRNA-seq data were passed on to several public bulk RNA-seq and proteome datasets to investigate cell type abundance using the cellular deconvolution approach in terms of COVID-19. Notably, two references, the modified LM22 (a gene signature matrix to distinguish 22 immune cell types) and the identified subtype (based on the scRNA-seq dataset) references, were used for cell type abundance analysis. Several predictors have been evaluated for their diagnostic performance in predicting severe disease. Our work holds great value for both clinical and public health for understanding the heterogeneity of neutrophils and other cell populations in multiple diseases.

**Figure 1.**
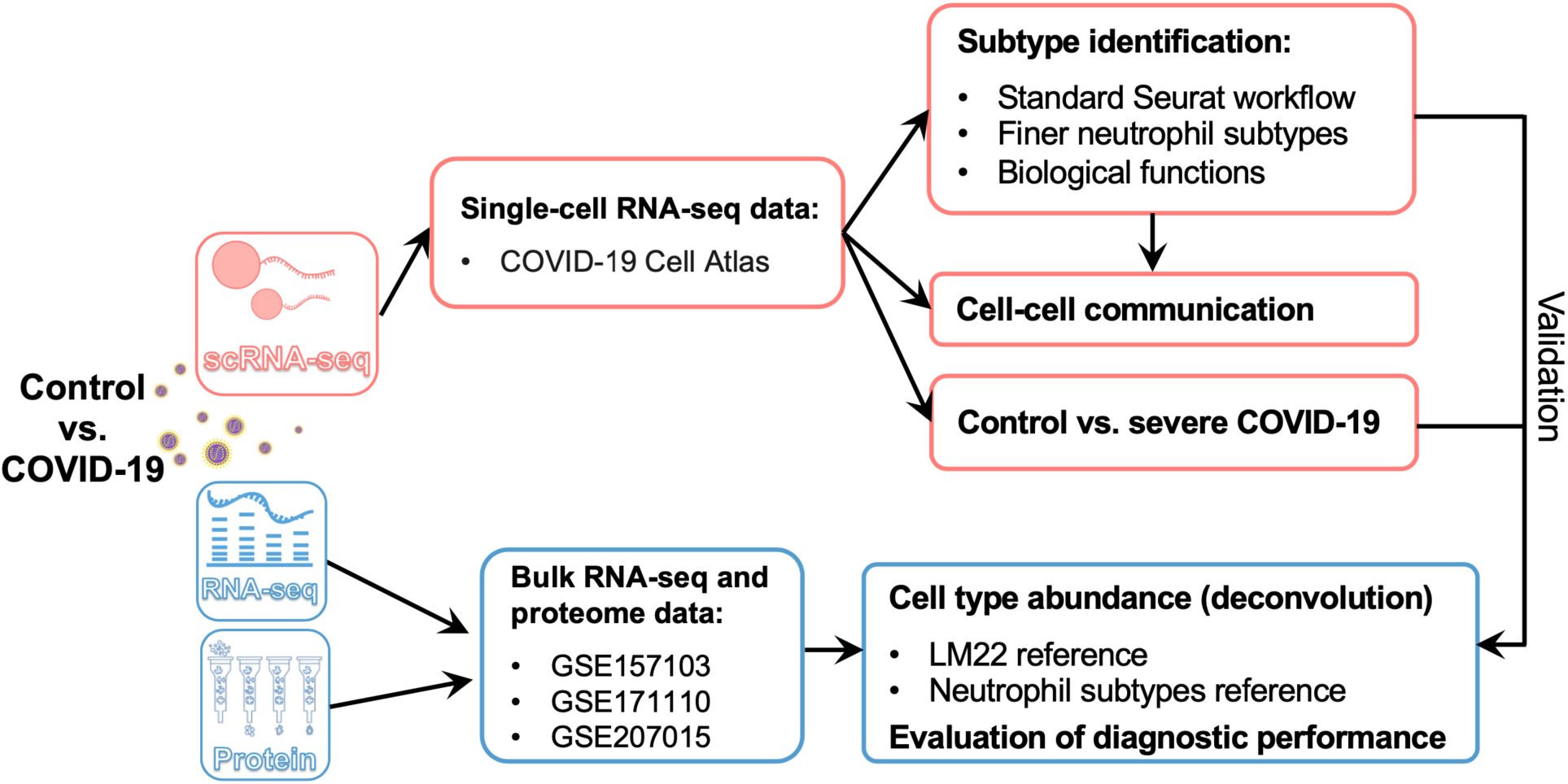
Workflow of this study. The scRNA-seq dataset was downloaded from the COVID-19 Cell Atlas. Neutrophils were used for further analyses, including finer subtype identification and cell–cell interactions. The disease and control groups were compared to investigate the mechanisms underlying disease pathogenesis. Finally, the results obtained using scRNA-seq data were validated using public bulk RNA-seq and proteome data.

## Material and Methods

### Data acquisition

Single-cell RNA sequencing (scRNA-seq) data were acquired from the COVID-19 Cell Atlas [11] and included healthy individuals and patients with different severity levels of COVID-19. The World Health Organization (WHO) score was used to evaluate disease severity as follows: healthy (WHO score 0), mild (WHO score 1–3), moderate (WHO score 4–5), severe (WHO score 6–7), and fatal (WHO score 8). scRNA-seq data were used to extract neutrophils and characterize the heterogeneity of their phenotypes and contributions to the disease. Two bulk RNA sequencing (bulk RNA-seq) datasets with accession numbers GSE157103 [7] and GSE171110 [30], and one human plasma proteome dataset (GSE207015) [31] were downloaded from the Gene Expression Omnibus (GEO) database to validate the identified neutrophil phenotypes in the scRNA-seq data and evaluate the performance of disease prediction models for severe COVID-19. For GSEA157103, the Sequential Organ Failure Assessment (SOFA) score was used to define the disease severity. Nine healthy individuals and 56 patients with COVID-19 were selected for the validation analysis. For the GSE171110 cohort, ten healthy and 44 severely ill samples were used. For the GSE207015 dataset, 41 healthy donors and 52 patients with different severities of COVID-19 were assessed. Additional details on the clinical and sample characteristics are presented in Additional file 7: Table S1.

### scRNA-seq data analysis and neutrophil subtype identification

Neutrophils were extracted from public blood scRNA-seq data, which included healthy individuals and patients with different severities of COVID-19 patients. Next, the obtained subset was processed using Seurat package [32] in R via the following steps: (i) Gene expression per cell were normalized based on its total expression using the “LogNormalize” method. (ii) The FindVariableFeatures function was used to Identify highly variable genes (HVGs), and their expression was scaled and centered using the ScaleData function. (iii) To perform dimensional reduction analysis, the scaled data and HVGs were passed to the RunPCA function for principal component analysis (PCA). (iv) The FindNeighbors and FindClusters functions were used to cluster cells by applying a graph-based clustering approach. Subsequently, clusters of these cells were visualized using a uniform manifold approximation and projection (UMAP) plot. Thirty principal components were used in this study. For the clustering analysis, the resolution parameter, which indirectly affects how many clusters can be identified, varied from 0.1 to 1, with an increment of 0.1 for each subsequent value. We then used Clustree [33] to visualize cell movement and detected clusters at multiple resolutions. The optimal resolution was jointly determined by further identification of markers and representative pathways. Eight clusters (neutrophil subtypes) were used for further analyses.

To identify signatures among the eight subtypes, the FindAllMarkers function was used with the following settings: min.pct = 0.25, logfc.threshold = 1, test.use = “wilcox” (a Wilcoxon Rank Sum test). Finally, the signatures were decided with additional filtering of adjusted P value ≤ 1 × 10^−10^. When comparing patients with severe COVID-19 to healthy individuals, FindMarkers function was used with min.pct = 0.25, test.use = “wilcox”, and logfc.threshold = 0.25. Then, the differentially expressed genes (DEGs) were selected with an additional cutoff of adjusted P value ≤ 0.1and |avg_log2FC| ≥ 1. The DEGs were visualized using a volcano plot. Differentially expressed transcription factors (TFs) were selected from the DEGs using a human TF list downloaded from AnimalTFDB v4.0 [34].

### Functional enrichment analysis

To describe the phenotypes and functions of neutrophil subtypes, we performed Gene Ontology (GO) and Kyoto Encyclopedia of Genes and Genomes (KEGG) pathway enrichment analyses using clusterProfiler [35]. GO enrichment analysis was performed based on biological process (BP), cellular component (CC), and molecular function (MF). Gene Set Enrichment Analysis (GSEA) was performed to uncover the mechanisms of COVID-19 pathogenesis using C7 immunologic signature gene sets from the Molecular Signatures Database (MSigDB; https://www.gsea-msigdb.org/gsea/msigdb). The enriched pathway was filtered by a cutoff P-value ≤ 0.05.

### Cell–cell communication analysis

CellChat [36] was used to explore cell–cell communications (CCC) among neutrophil subtypes and other cell types. A secreted signaling pathway database, including ligand–receptor (L–R) pairs, was used to infer cellular interactions. The label of neutrophil subtypes per cell was passed to the corresponding cells in the original scRNA-seq data to investigate cell–cell interactions among/within neutrophil subtypes and other cell types. Biologically significant signaling changes were selected to visualize communication patterns and networks.

### Cellular deconvolution and disease predictions using bulk RNA-seq and proteome data

CIBERSORTx [37] approach was used to estimate the proportion of cell types in the bulk RNA-seq and proteome datasets. Two references were used for this analysis: modified public expression signatures (LM22) and the signatures of neutrophil subtypes. The original LM22 reference data, provided by CIBERSORTx, included the signature expression of 22 mature human hematopoietic populations from blood. We modified this reference by merging subtypes or similar types of cells into 12 main cell types using an average value. For the customized neutrophil subtype reference, the average expression of the detected signatures was calculated. In addition, receiver operating characteristic (ROC) curves were used to evaluate the performance of several predictors of severe COVID-19.

### Cell senescence status analysis

Multiple factors, including oncogene activation, DNA damage, tissue damage, aging, and viral infections, can trigger senescence [38,39]. Thus, to study the cell senescence status of neutrophil subtypes, three gene sets were used. One of these was human aging gene set, which included 307 genes downloaded from the aging gene database (https://genomics.senescence.info/genes/human.html). The other two gene sets were acquired from the MsigDB, which were gene sets of Fridman senescence UP (77 genes) [40] and Reactome cellular senescence (214 genes). The cell senescence status of neutrophil subtypes was evaluated according to the following steps: (1) for a cell, the average expression of all genes in a senescence gene set was calculated; (2) based on the above average values, all cells from a subtype were ranked, and then the top 10% of cells were defined as senescent cells; (3) for each subtype, the relative senescent cell percentage was calculated by dividing the number of senescent cells (top 10% of cells in a subtype) by the total cell number in this subtype. The percentage of senescent cells was used to evaluate the cellular senescence status of the eight neutrophil subtypes.

### Statistical analysis

Two-sample variables were estimated using the non-parametric Wilcoxon rank-sum test. Significant levels are represented in the violin plot as p < 0.05, < 0.01, and < 0.001. All analyses were conducted using the R (version 4.1.1).

## Results

### Heterogeneous neutrophils detected in scRNA-seq data

scRNA-seq data obtained from blood samples of healthy individuals and patients with COVID-19[11] revealed a total of 174,753 cells that were observed in the original study; these were categorized into 15 cell types, including the neutrophil population (Additional file 1: Figure S1). Subsequently, a subset of 39,845 neutrophils was extracted to study their phenotypic and functional heterogeneity; this subset was used for further analyses in the present study. Based on the embedding dimension reduction of the source data, we visualized the obtained neutrophils on a UMAP plot in terms of disease severity and observed that some neutrophil subpopulations showed different abundances in terms of COVID-19 severity, suggesting that neutrophils are heterogeneous and that phenotypes with different functions may exist under physiological and pathological conditions (Additional file 1: Figure S1).

### Identification of neutrophil subpopulations and their enriched pathways

Using scRNA-seq data of neutrophils, we identified finer subtypes of neutrophils and described their heterogeneous functions. First, the standard Seurat workflow for scRNA-seq data processing was used to identify finer neutrophil subtypes. Combining this analysis with the visualization of cell movements at multiple resolutions that were used for clustering (Additional file 2: Figure S2), eight neutrophil subtypes, namely C1–C8, were determined based on the selected resolution (Figure 2A). We calculated the compositions of the eight identified subtypes in terms of COVID-19 severity (Figure 2B–D); we found that the C1 and C8 subtypes showed the highest lowest fractions, respectively, among the neutrophil subtypes. Moreover, the neutrophil subtype composition varied according to the disease severity. Specifically, the C1 subtype exhibited the highest fraction in patients with mild, moderate, and severe disease, whereas the C2, C3, and C5 subtypes were abundant in healthy individuals. The C4 subtype was particularly prevalent among severely ill and fatal donors. The C6, C7, and C8 subtypes exhibited some degree of extensions in patients with COVID-19. Next, the activation signatures of the eight neutrophil subtypes were detected; several subtype-specific genes among the signatures were considered as markers for the subtypes (Additional file 8: Table S2, Figure 2E). The neutrophil subtypes were renamed by adding the subtype-specific gene symbols IFIT1/2/3 (C1), IL7R (C2), HSPA5 (C3), DEFA1/1B/3 (C4), G0S2 (C5), FKBP5 (C6), MMP8 (C7), and HBB (C8) (Figure 2A). Furthermore, the number of signatures varied considerably. Compared with most of the C7_MMP8 (n=137) and C8_HBB (n=153) genes, only a few signatures were detected in the C3_HSPA5 (n=3) and C6_FKBP5 (n=16) subtypes (Figure 3A). Next, we visualized the expression of the signatures and performed GO and KEGG pathway enrichment analyses to characterize the functions of each subtype (Figure 3B–H). Notably, the enriched pathways of C3_HSPA5 and C6_FKBP5 were not detected because of the small number of signatures; thus, their related functions could not be described. We found that the subtypes displayed unique or commonly enriched pathways, which may lead to different phenotypes or contribute to different conditions (e.g., healthy and COVID-19). Specifically, C2_IL7R, C5_G0S2, and C8_HBB were associated with the ribosome pathway and transcription activator activity, while C1_IFIT1/2/3, C4_DEFA1/1B/3, and C7_MMP8 were associated with neutrophil-mediated immunity, glycolysis/gluconeogenesis, NOD-like receptor signaling pathway, and HIF-1 signaling pathway.

**Figure 2.**
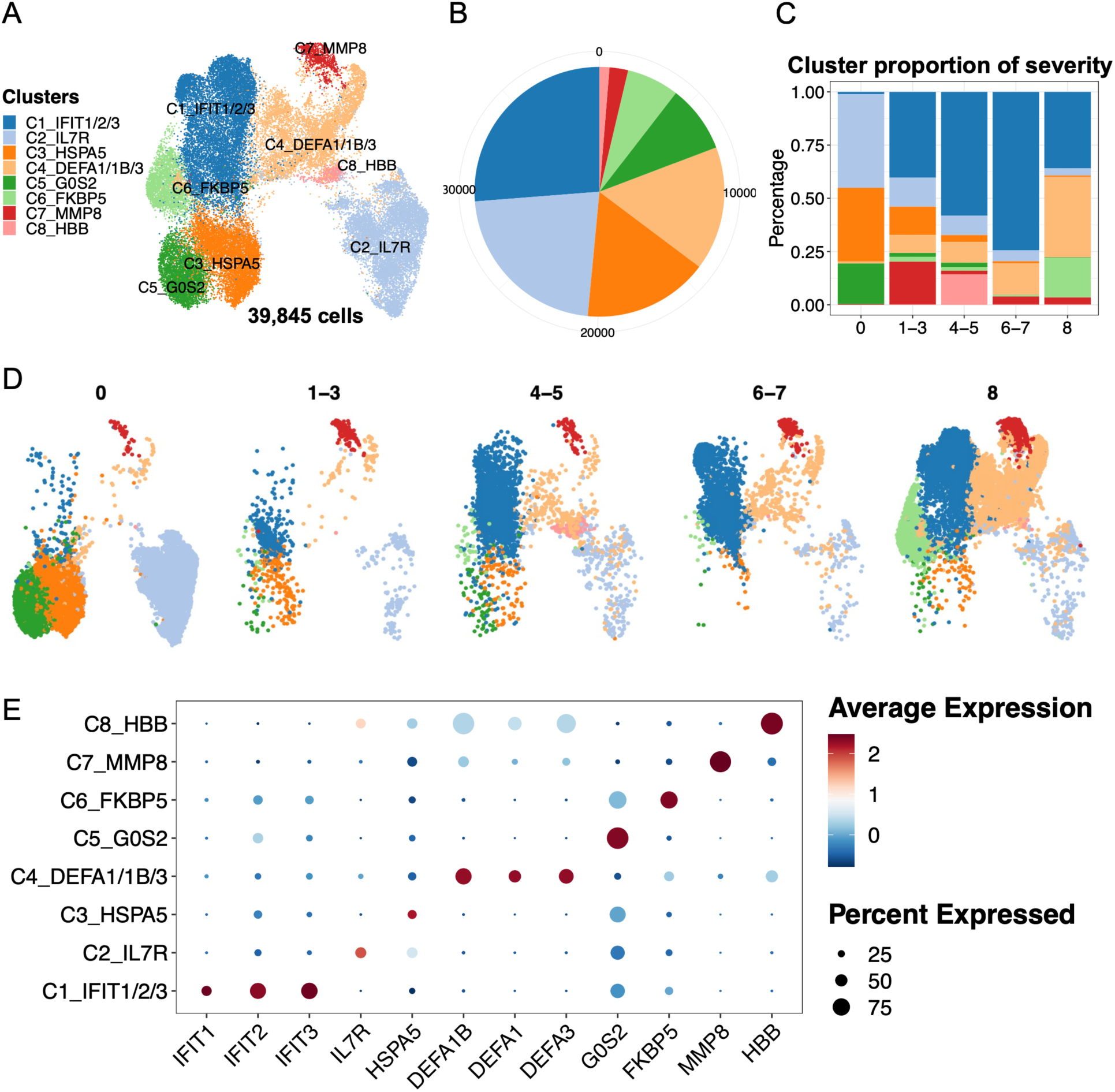
Characterization of neutrophil heterogeneity. (A) The UMAP plot exhibits the eight identified neutrophil subtypes. (B) The pie plot indicates the cell numbers in each subtype. (C) The bar plot shows subtype compositions across the severity of COVID-19. (D) The UMAP plots display the subtypes in terms of COVID-19 severity. (E) The dot plot represents the selected markers for each subtype. The WHO scores were used to classify healthy (WHO score 0), mild (WHO score 1–3), moderate (WHO score 4–5), severe (WHO score 6–7), and fatal (WHO score 8) donors.

**Figure 3.**
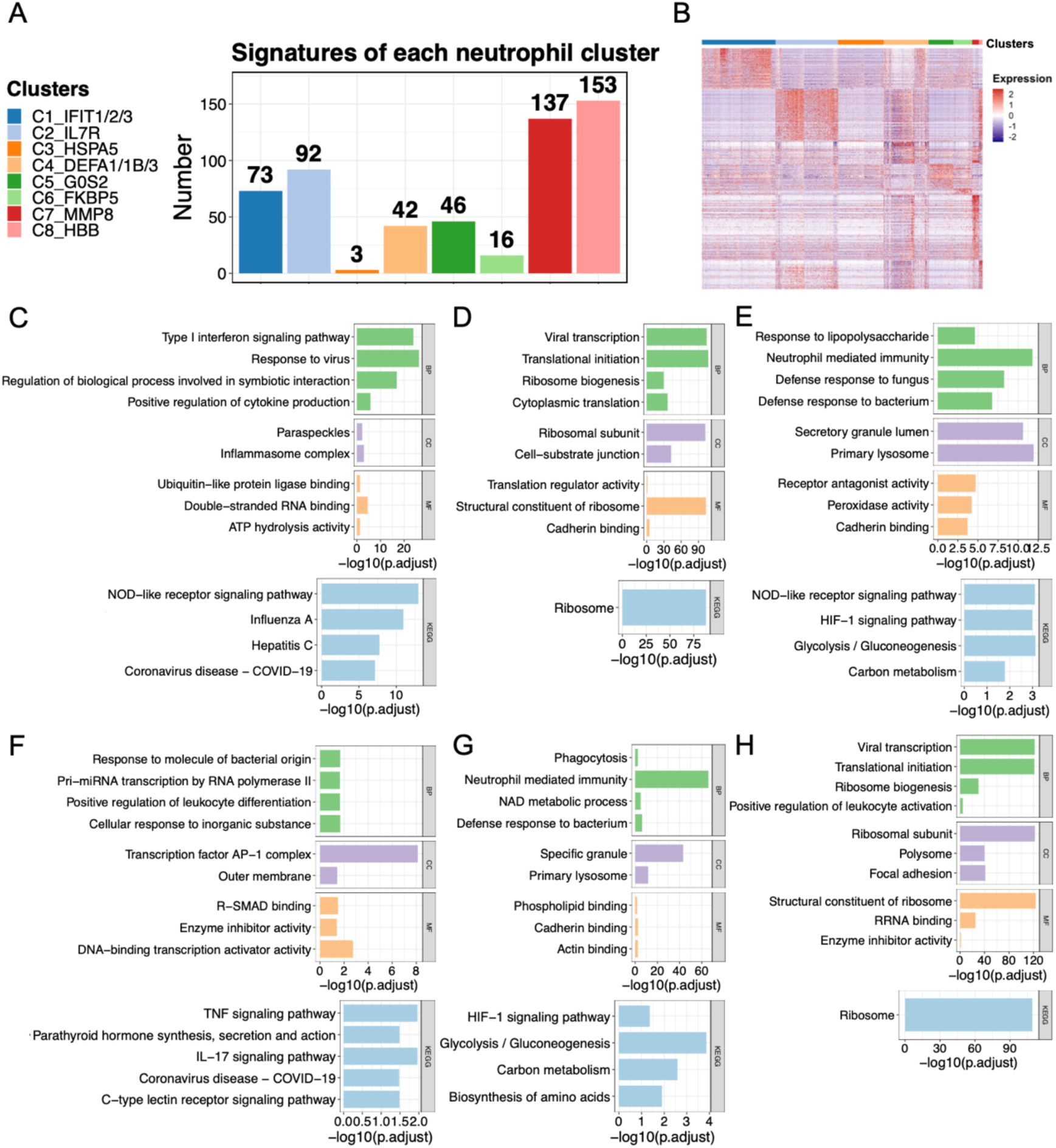
Biological functions of each neutrophil subtype. (A) Gene signatures of the eight subtypes. (B) Heatmap represents the signature expression in (A). The identified signatures in (A) were used for enrichment analysis. (C–H) The representative enriched GO and KEGG pathways of each subtype are presented: (C) C1_IFIT1/2/3, (D) C2_IL7R, (E) C4_DEFA1/1B/3, (F) C5_G0S2, (G) C7_MMP8, and (H) C8_HBB. The representative pathways of C3_HSPA5 and C6_FKBP5 are not provided owing to the small size of identified signatures. GO enrichment analysis was performed based on BP, CC, and MF. GO, Gene Ontology; KEGG, Kyoto Encyclopedia of Genes and Genomes; BP, biological process; CC, cellular component; MF, molecular function.

### Neutrophil phenotypes can be altered under physiological and pathological conditions

The phenotypes and functions of the eight identified neutrophil subtypes were further described by studying their cell interactions among the subtypes and with other cell types. To understand how neutrophil subtypes interact with other cell types such as monocytes, B cells, and T cells, we analyzed CCC using the CellChat tool to identify representative L–R pairs. The eight subtype labels (C1–C8) were transferred to the original scRNA-seq data, which included other cell types and neutrophils. We inferred the signaling pathways and communication patterns and then visualized the representative signaling pathways in the networks (Figure 4). Five outgoing and incoming communication patterns were identified between cell types and their associated signaling pathways. Outgoing communication patterns indicate how secreting cells (acting as senders) drive interactions via specific signaling pathways by secreting ligands. In contrast, incoming communication patterns demonstrate how target cells (acting as receivers) contribute to interactions by expressing their receptors. Regarding the results of both the outcoming and incoming patterns in terms of cell types, we found that neutrophil subtypes could be classified into various patterns connected to eight representative signaling pathways. This suggests that the identified pathways can have different roles as senders or receivers depending on the subtypes to which they pertain. To study in detail how each subtype interacted with others in a specific pathway, we visualized six of the identified signaling pathways in the networks (Figure 4C). Two pathways, the CCL and PARs signaling pathways, were excluded because cell interactions were only detected among NK cells, monocytes, and T cells, rather than neutrophils. We found that the roles of neutrophil subtypes may vary for each signaling pathway. For instance, all eight subtypes received signals from monocytes or dendritic cells in the GALECTIN signaling pathway, whereas only the C7_MMP8 subtype sent signals to all other cell types in the RESISTIN signaling pathway.

**Figure 4.**
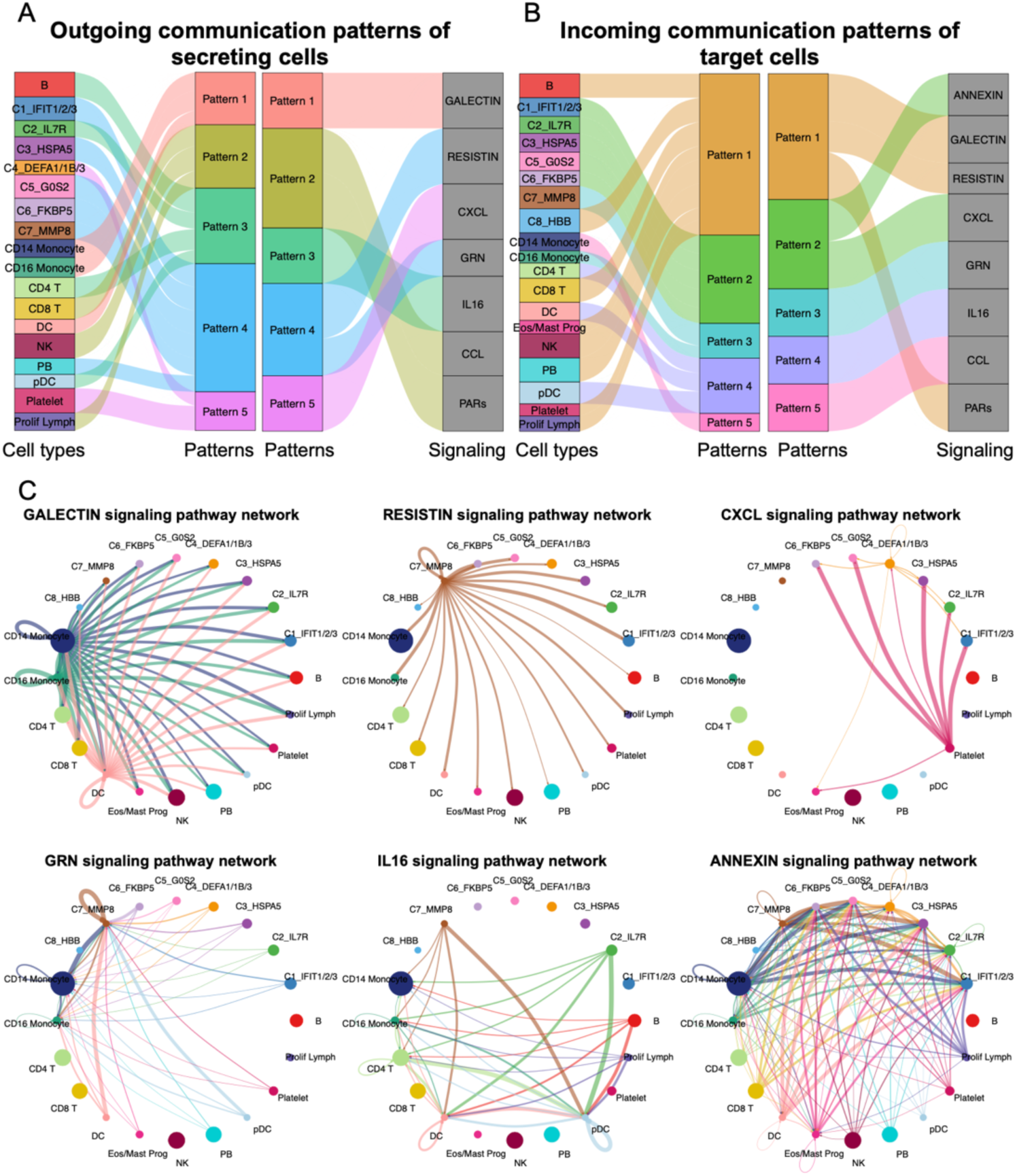
Cell–cell communications among the eight identified neutrophil subtypes and other cell types. Cell–cell communications (CCCs) among neutrophil subtypes and other cell types was performed by focusing on ligand–receptor (L–R) pairs. The (A) outgoing and (B) incoming communication patterns are represented for each cell type. Alluvial plots were used to visualize the cell types, inferred latent patterns, and associated signaling pathways. Flow thickness refers to the degree of contribution of cell types or signaling pathways to the latent patterns. (C) Networks indicate the CCCs of representative signaling pathways between neutrophil subtypes and others. Cell types are represented by different colors. The size of the circle is proportional to the cell number. The edge width represents the number of L–R pairs. Arrows represent the direction of interactions from one cell (sender) to another (receiver).

We created a diagram showing L–R pairs in the six signaling pathways and their expression among the neutrophil subtypes (Figure 5 and Additional file 3: Figure S3). The diagram reveals five ligands (RETN, GRN, IL16, ANXA1, and PPBP) and five receptors (SORT1, CXCR2, CD45, CAP1, and FPR1). The C6_FKBP5 subtype expressed five receptors and no ligands, whereas the C7_MMP8 subtype expressed four ligands and two receptors. No ligands were detected in the C1_IFIT1/2/3, C3_HSPA5, C5_G0S2, or C8_HBB subtypes. In C7_MMP8, SORT1 and RETN were more highly expressed than in other subtypes (Figure 5B). Interestingly, the PPBP ligand (pro-platelet basic protein) of the CXC chemokine family was highly expressed in C4_DEFA1/1B/3, which was associated with severe and fatal COVID-19 cases. Notably, PPBP was an activation signature of the C4_DEFA1/1B/3 subtype (Figure 5B and Additional file 8: Table S2). A previous study on PPBP reported that a subpopulation of neutrophils with high PPBP level may correspond to neutrophil–platelet aggregates and, therefore, could serve as a promising blood-based prognostic biomarker for solid tumors, offering clinical value in their assessment [41]. Our findings demonstrate that alterations in neutrophil phenotype can affect disease pathogenesis. Therefore, PPBP may be a useful biomarker for the diagnosis of severe COVID-19.

**Figure 5.**
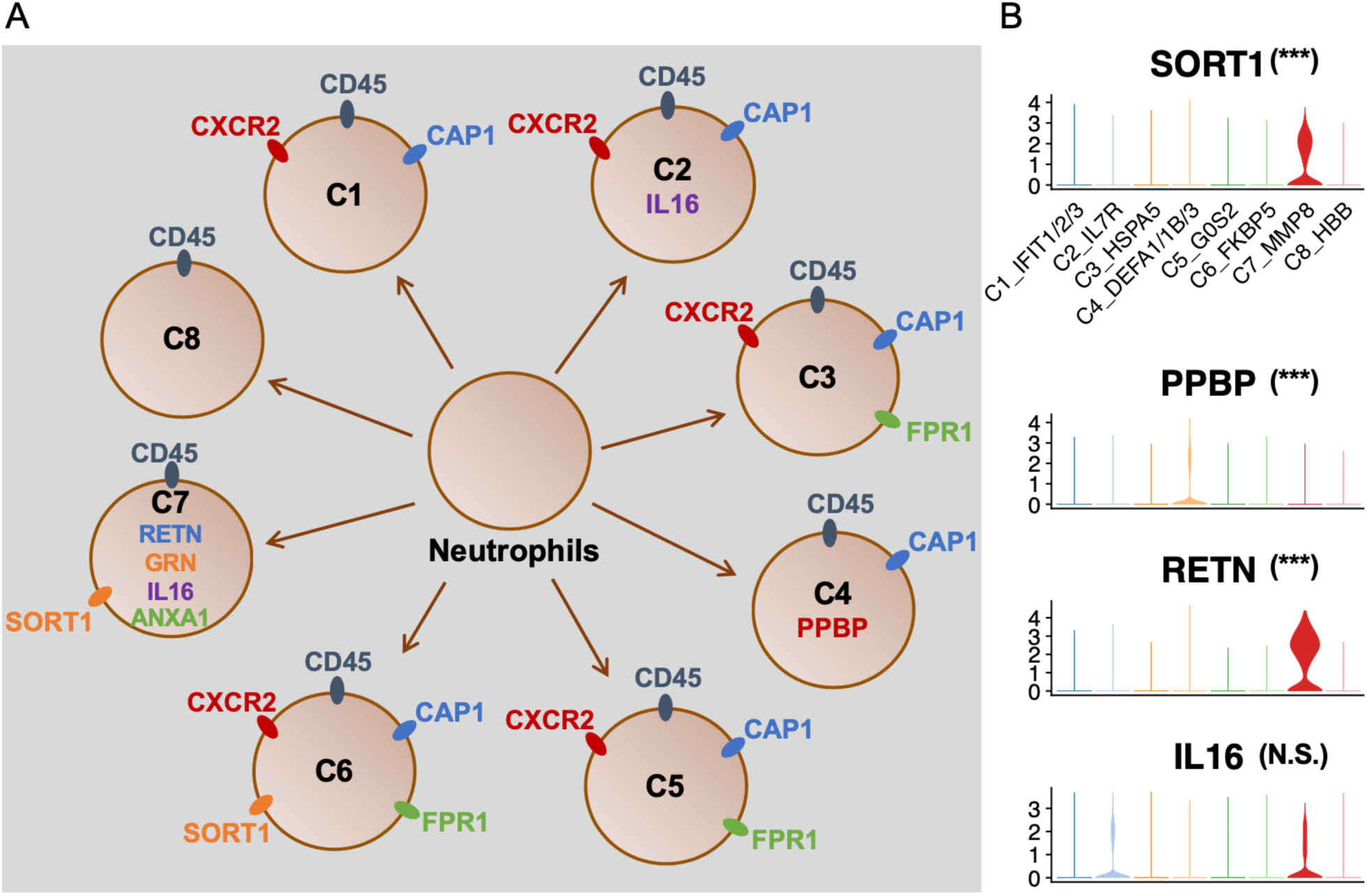
Schematic diagram for the neutrophil phenotypes. (A) Schematic diagram for the eight identified neutrophil subtypes. Ligands and receptors for each subtype are shown inside and outside the cell, respectively. The same color marks the shared ligands and receptors among subtypes. (B) Violin plots represent expression of selected ligands and receptors. Significant gene expression was determined by comparing a relative subtype to all the remaining subtypes. * p-value < 0.05; ** p-value < 0.01; *** p-value < 0.001; N.S., non-significant.

### Severe COVID-19-specific expression patterns in neutrophils

Then, we compared the neutrophils of healthy donors with that of patients to reveal the mechanisms underlying COVID-19 progression. Specifically, we identified DEGs between severe COVID-19 and healthy cells; a total of 55 upregulated and 38 downregulated genes were identified (Additional file 9: Table S3), which are marked in red and blue, respectively, in the volcano plot (Figure 6). We also investigated enriched pathways to uncover the mechanisms underlying disease progression (Figure 6). GO and KEGG pathway enrichment analyses demonstrated that the upregulated genes were associated with influenza response and certain signaling pathways, such as the NOD-like receptor, RIG-I-like receptor, IL-17, and Toll-like receptor signaling pathways. In contrast, the downregulated genes were related to ribosome pathways. Several studies have demonstrated the important roles of these signaling pathways in either innate immunity or COVID-19 progression, including their utility as diagnostic biomarkers and potential treatment strategies [42–48]. In addition, we found that some TFs were differentially expressed in severe COVID-19. Specifically, IRF7, SP100, HMGB2, and BCL6 TFs were upregulated, whereas FOS and JUNB were downregulated. We also performed GSEA based on the transcriptome of patients with severe disease and found that 971 gene sets were enriched with a P-value cutoff of 0.05 (Additional file 10: Table S4). We visualized three representative enriched gene sets and marked multiple S100 family members (e.g., S100A8/A9), interferon (e.g., IFITM2), and MMP family members (e.g., MMP8), as well as cytokines, chemokines, and their receptors (e.g., IL1R2 and CXCR1) (Figure 6D). The selected genes showed high expression levels in the severe COVID-19 transcriptome.

**Figure 6.**
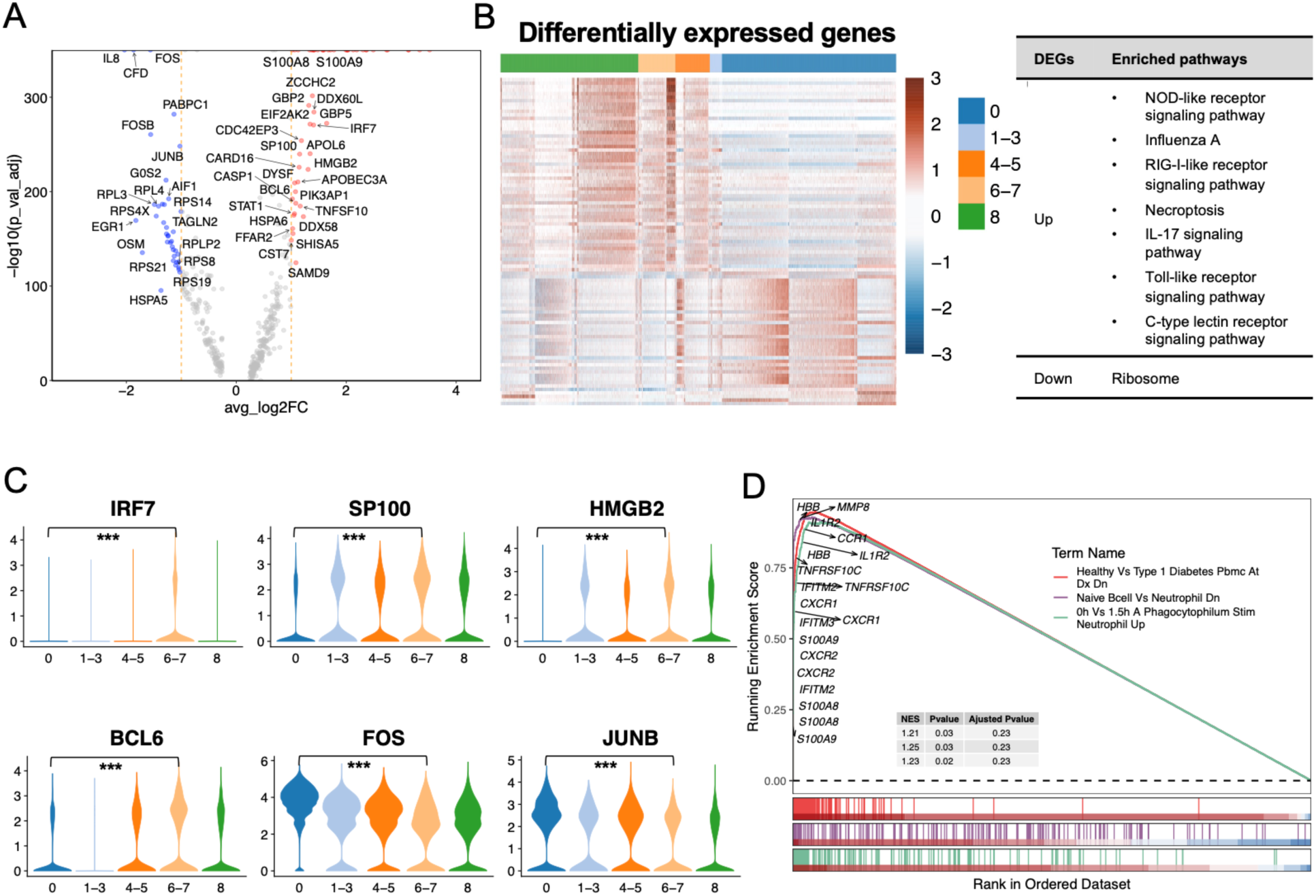
Comparisons of healthy individuals and patients with severe COVID-19. (A) Volcano plot represents the DEGs in patients with severe COVID-19 compared to healthy individuals. Red, blue, and grey colors represent upregulated, downregulated, and stable genes in severe COVID-19 cases, respectively. (B) Heatmap shows the DEGs in severe COVID-19 when comparing to healthy individuals. Five WHO severity score groups: healthy (0), mild (1–3), moderate (4–5), severe (6–7), and fatal (8). (C) Selected differentially expressed TFs. Significant gene expression was determined by comparing the severe (6–7) and healthy (0) groups. * p value < 0.05; ** p value < 0.01; *** p value < 0.001. (D) GSEA analysis was performed based on the C7_MMP8 immunologic signature gene sets from MSigDB. GSEA: gene set enrichment analysis.

Recent studies have identified virus-induced senescence is the underlying cause of cytokine release and inflammation in patients with COVID-19, and a robust upregulation of the senescence phenotype in the majority of COVID-19 patients was particularly observed in lung epithelial cells after viral infection [39,49,50]. Considering these studies, we investigated cell senescence status across the eight neutrophil subtypes using three senescence-associated gene sets (Additional file 4: Figure S4). Some of the neutrophil subtypes abundant in COVID-19 (e.g., C2_IL7R, C6_FKBP5, and C7_MMP8) showed high levels of senescent cells, whereas others (C4_DEFA1/1B/3 and C8_HBB) exhibited low levels. Our findings indicate that cellular senescence may not be commonly triggered in SARS-CoV-2-infected neutrophils compared to that in infected lung epithelial cells, and that senescence induction is dependent on the subpopulations of neutrophils.

### Validation of C4_DEFA1/1B/3 neutrophil subtype abundance as predictor of COVID-19 severity

After investigating the phenotypes and functions of the identified neutrophil subtypes and the mechanisms of COVID-19, a validation analysis was performed. We used two bulk RNA-seq datasets and one human plasma proteome dataset (accession numbers GSE157103, GSE171110, and GSE207015) to validate the neutrophil subtypes identified in the scRNA-seq data and thus evaluate the performance of severe COVID-19 prediction models. On the one hand, cellular deconvolution analysis using a modified LM22 expression signature reference indicated that the relative abundances of neutrophil subpopulations were increased in severe COVID-19 patients for the two bulk RNA-seq and proteome datasets (Additional file 5: Figure S5A). On the other hand, the signature matrix of identified neutrophil subtypes was used as a customized reference to investigate their relative abundances. We observed that the C4_DEFA1/1B/3 fractions, which showed an expansion in severe and fatal cases in the scRNA-seq data, increased along with the SOFA score or disease severity (Figure 7). In addition, we calculated the Pearson correlation between the fraction of each neutrophil subtype and the SOFA score using the GSE157103 dataset and found that some neutrophil subtypes, such as C4_DEFA1/1B/3, were positively correlated with the SOFA score (Additional file 5: Figure S5B).

**Figure 7.**
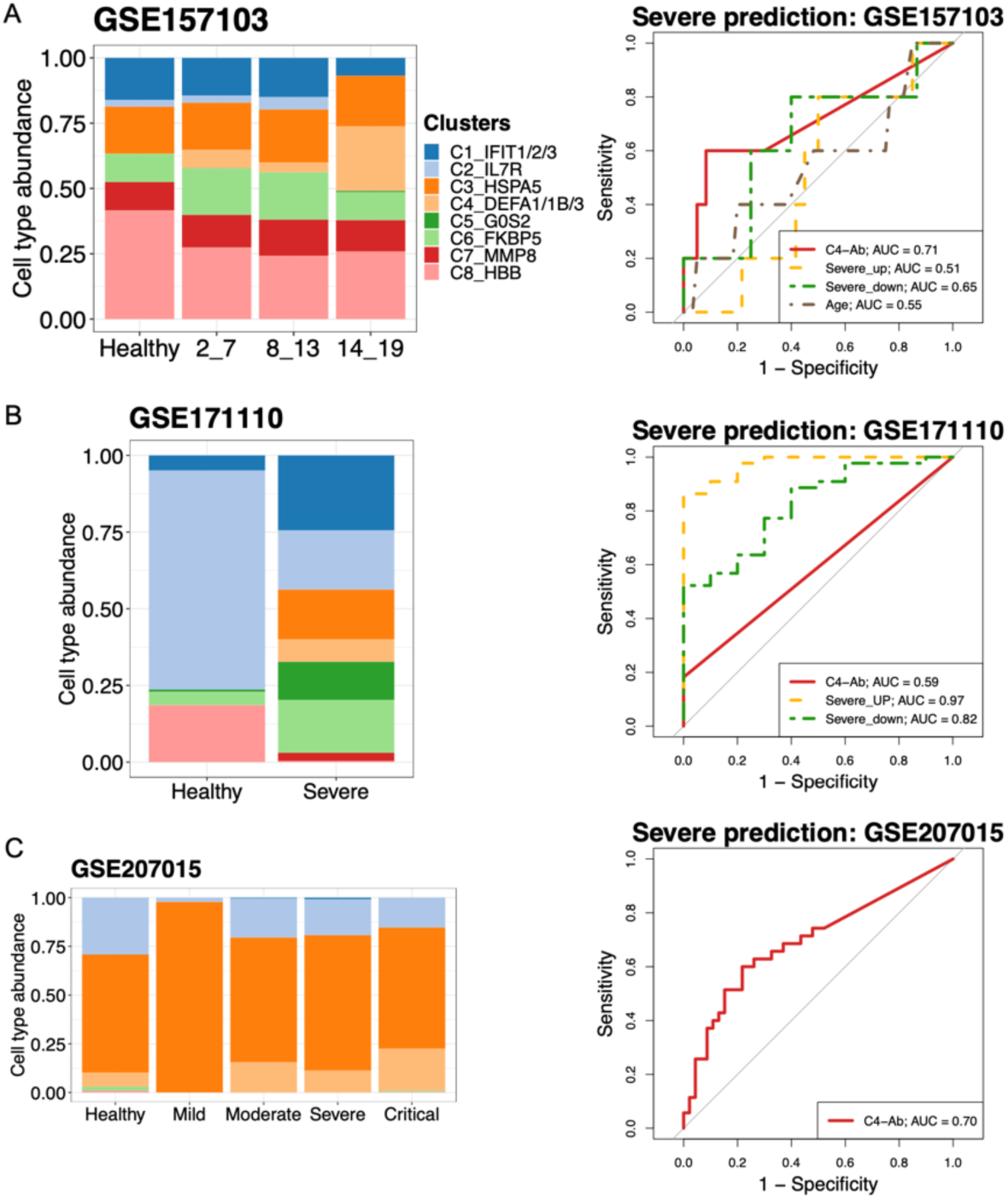
Cell type abundance estimation and severe predictions using bulk RNA-seq and human plasma proteome datasets. (A–C) Analyses performed using GSE157103, GSE171110, and GSE207015, respectively. Cellular deconvolution analysis was performed using the eight neutrophil subtypes identified in the scRNA-seq dataset. The bar plots indicate the eight subtype proportions between healthy individuals and patients with COVID-19. The average subtype fractions for control–case groups were used. In the case of GSE157103, SOFA scores varying from 2 to 19 were used to evaluate disease severity. For GSE171110, healthy and severely ill patients were analyzed. For GSE207015, the COVID-19 severity was classified as mild, moderate, severe, and critical. The right panels represent disease predictions for multiple predictors using the area under the receiver operating characteristic (ROC) curve (AUC). Predictors included C4-Ab (abundance of neutrophil subtype C4_DEFA1/1B/3), specific differentially expressed gene sets in severe cases, and/or based on age; upregulated genes: IRF7, SP100, HMGB2, and BCL6; downregulated genes: FOS and JUNB. The relative expression levels of these genes in the scRNA-seq dataset are shown in Figure 6D. SOFA: sequential organ failure assessment.

Considering the abundance of the C4_DEFA1/1B/3 fraction in patients with severe disease, we used area under the receiver operating characteristic curve (AUC) analysis to evaluate the diagnostic performance for severe COVID-19 prediction using various predictors, including the abundance of neutrophil C4_DEFA1/1B/3 subtype (C4-Ab), and identified differentially expressed TFs (upregulated: IRF7, SP100, HMGB2, and BCL6; downregulated: FOS and JUNB). Notably, the differentially expressed TF combinations were not used for prediction in the proteome dataset, as some proteins were not observed in the proteome matrix. In the GSE157103 dataset, the C4_DEFA1/1B/3 fraction showed the highest performance in predicting severe COVID-19, whereas the upregulated TF sets displayed remarkable performance in the GSE171110 dataset. Collectively, the percentage of C4_DEFA1/1B/3 neutrophil subtypes exhibited good or moderate diagnostic performance for distinguishing severe COVID-19 from healthy status in the two-validation bulk RNA-seq and one proteome datasets (AUC = 0.71, AUC = 0.59, and AUC = 0.7, respectively). The upregulated TFs showed excellent diagnostic value in the GSE171110 (AUC = 0.97).

## Discussion

The SARS-CoV-2 virus is responsible for COVID-19, an extremely contagious respiratory infection. Several studies have shown that neutrophils are activated in response to viral infections. Neutrophils play a critical role in the immune system and can help fight infections and inflammation-induced injuries. Moreover, neutrophil phenotypes are considered to be altered and plastic in response to the pathogenesis and progression of many diseases. For example, some subpopulations are associated with an unfavorable cancer prognosis or the release of specific cytokines that inhibit inflammation [12,18,21]. Studies on the phenotypic and functional diversities of neutrophils help us understand the role of neutrophils in antimicrobial functions and pathogenic immune processes. However, the functional descriptions of the various neutrophil phenotypes after SARS-CoV-2 infection are still not fully understood.

This study identified and validated eight neutrophil subtypes that showed either common or diverse functions by analyzing scRNA-seq, bulk RNA-seq, and proteomic datasets. The identified neutrophil subtypes were characterized by distinct features, including various numbers of activated signatures, different representative signaling pathways, and multiple enriched molecular functions. Some subtypes related to healthy cells possess multiple functions, such as enriched ribosome biosynthesis and transcription/translation activity. Some genes that were associated with virus-infected cells showed significant enrichment in immune responses, metabolic processes, and various signaling pathways. These findings suggest that subsets of neutrophils show various phenotypic and functional characteristics and that such a wide variety of neutrophil subpopulations may lead to diverse responses to diseases. For example, C4_DEFA1/1B/3 neutrophils are associated with severe and fatal diseases. Numerous activation signatures (42 genes) were detected, showing considerable enrichment in neutrophil-mediated immunity and some signaling pathways. CCC analysis further indicated the presence of a representative CXCL signaling pathway (i.e., the L–R pair of PPBP and CXCR2) in the C4_DEFA1/1B/3 population. The PPBP ligand, one of the signatures of the C4_DEFA1/1B/3 subtype in this study, is involved in various conditions; PPBP was detected in activated platelets that are involved in chronic inflammation during the development of atherogenesis and coronary heart disease (CHD), and thus could be used as a biomarker for CHD risk in postmenopausal Thai women [51]. Combined with PADI4 expression, PPBP expression may be considered a non-invasive multi-marker approach offering diagnostic and clinical value in patients with suspected lung cancer [51]. Another study demonstrated that PPBP is a major repressive target of glucocorticoids that promote gluconeogenesis [52,53]. Interestingly, we found that the glycolysis/gluconeogenesis pathway was enriched in the C4_DEFA1/1B/3 subtype (Figure 3E). Additionally, transmembrane receptor expression of CD45 and CAP1 was detected in the C4_DEFA1/1B/3 fraction. Our phenotypic and functional descriptions of the C4_DEFA1/1B/3 subtype may provide insights into the specific responses in severe and fatal cases. Significant expression of PPBP in the C4_DEFA1/1B/3 subtype may be a novel biomarker of severe COVID-19.

However, owing to the small size of the activation signatures, our study failed to describe the phenotypic and functional features of the C3_HSPA5 and C6_FKBP5 subtypes. Compared to the smallest total number of counts and genes across the eight neutrophil subtypes, the values of the C6_FKBP5 subtype were intermediate (Additional file 6: Figure S6). Together with the relatively large number of cells of these two subtypes (Figure 2A, B), we hypothesized that C3_HSPA5 and C6_FKBP5 may be fundamental populations, that share most features with other neutrophil subpopulations. In addition, we obtained discrepant results regarding the correlation between neutrophil subtype fractions and SOFA scores. For example, the C8_HBB population was found to be activated post-viral infection, and many signatures were observed (137 genes) using scRNA-seq data; however, this fraction negatively correlated with the SOFA score using bulk RNA-seq data (Figure 2A and B, Figure 3A, and Additional file 5: Figure S5). In addition, some neutrophil subtype abundances exhibited differences between transcriptome and proteome profiling, such as the highly abundant C3_HSPA5 subtype in the proteome (Figure 7). Therefore, increasing the size and type of datasets may help better interpret such results in the future. Moreover, future analyses, in combination with assessment of other diseases, may provide a comprehensive understanding of the heterogeneity of neutrophils and alterations in their phenotypes and functions.

## Conclusion

Collectively, our study identified several subpopulations of neutrophils in patients with SARS-CoV-2 infection by analyzing scRNA-seq, bulk RNA-seq, and proteome datasets and characterized the alterations in their phenotypes and functions from various aspects (e.g., signature identification, representative pathways, cellular components, and cell-cell interactions). Additionally, we found that PPBP may serve as a novel biomarker for diagnosing severe COVID-19 and may offer clinical value. Our study provides a framework for understanding the plasticity of neutrophils and other cell populations, and how a wide range of subpopulations respond to different diseases.

## Ethics approval and consent to participate

Not applicable.

## Consent for publication

Not applicable.

## Availability of data and materials

All the datasets could be downloaded from the indicated databases/websites. The names of the databases/websites and accession numbers (s) can be found in the article/supplementary materials.

## Competing interests

The authors declare that they have no competing interests.

## Funding

This research was partially supported by the Research Support Project for Life Science and Drug Discovery (Basis for Supporting Innovative Drug Discovery and Life Science Research (BINDS)) from AMED under Grant Number JP22ama121019; and by the project “Development of the Key Technologies for the Next-Generation Artificial Intelligence/Robots” of the New Energy and Industrial Technology Development Organization.

## Author contributions

LZ conceived and designed the study. LZ contributed to the data collection and analysis. HN and KK provided comments and suggestions on the methodology. LZ, HN and KK interpreted the results. LZ wrote the manuscript. HN and KK reviewed and edited the manuscript. All the authors contributed to the manuscript and approved the submitted version.

## Supplementary information

**Additional file 1: Figure S1. Characterization of the source dataset.**

The cells from the source scRNA-seq dataset were visualized using a UMAP plot. Fifteen cell types were identified in this study. A subset of cells (neutrophils only) was extracted and visualized according to COVID-19 severity. Five severity groups were defined as follows: healthy (0), mild (1–3), moderate (4–5), severe (6–7), and fatal (8).

**Additional file 2: Figure S2. Visualization of cell movement in terms of multiple resolutions.** Clustree was used to visualize and define the optimal resolution for cell clustering. For clustering, the resolution parameter varied from 0.1 to 1, with an increase of 0.1 for every step.

**Additional file 3: Figure S3. Expressions of selected ligands and receptors.**

Violin plots represent the expression of selected ligands and receptors, as shown in Figure 5. Significant gene expression analysis was performed by comparing a subtype with all other subtypes. * p-value < 0.05; ** p-value < 0.01; *** p value < 0.001; N.S., non-significant.

**Additional file 4: Figure S4. Cell senescence status across the eight neutrophil subtypes.**

The percentage of senescent cells in each cluster was calculated. Senescent cells were defined based on the average expression of the three gene sets (i.e., aging, Fridman, and reactome). The top 10% of cells in a given cluster were defined as senescent based on average expression levels.

**Additional file 5: Figure S5. Immune cell type abundance and correlation between neutrophil subtypes and SOFA score.**

(A) Cell type abundance estimation for GSE157103, GSE171110, and GSE207015 datasets using modified LM22 reference. (B) Pearson correlation between the eight neutrophil subtype fractions and SOFA scores using GSE157103 dataset. SOFA: sequential organ failure assessment.

**Additional file 6: Figure S6. Statistics of the eight characterized subtypes of neutrophils.**

For each cell, the box plots indicate the total number of (A) UMI counts and (B) genes.

**Additional file 7: Table S1. Clinical and sample characteristics of patients with COVID-19 and healthy donors.**

**Additional file 8: Table S2. Signatures of the eight neutrophil subtypes identified.**

**Additional file 9: Table S3. Differentially-expressed genes in patients with severe COVID-19 and healthy individuals.**

**Additional file 10: Table S4. Gene set enrichment analysis (GSEA) analysis for severe COVID-19 based on the C7 gene sets from MsigDB.**

## Notes

### Competing Interest Statement

The authors have declared no competing interest.

