## Supplementary material for "Multi-omics profiling reveals phenotypic and functional heterogeneity of neutrophils in COVID-19": Supplementary Figures.pdf

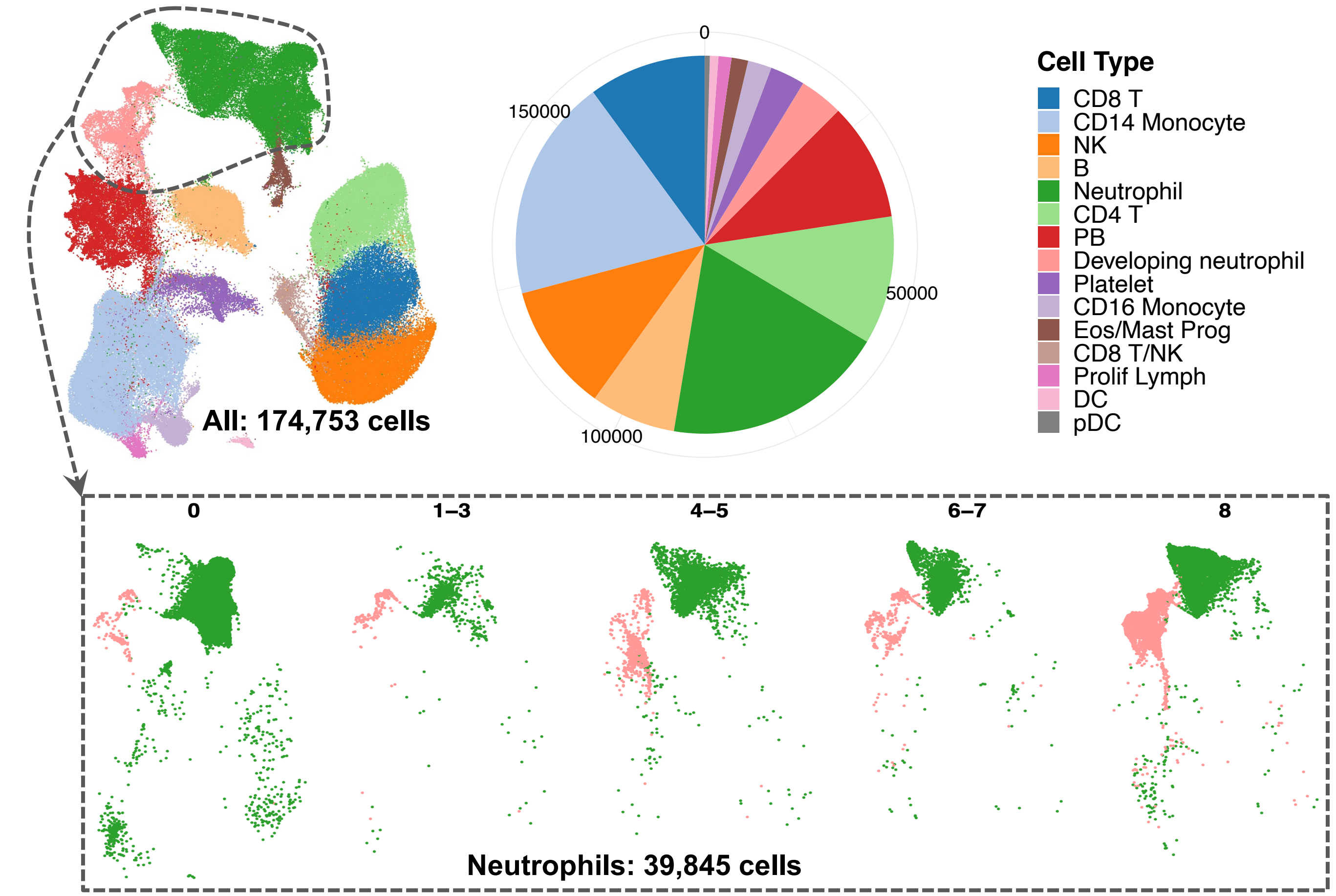


**Supplementary Figure 1. Characterization of the source dataset.**
Cells from the source scRNA-seq dataset were visualized on a UMAP plot. A total of 15 cell types were annotated. A subset of cells (neutrophils only) were extracted and visualized subpopulations in terms of COVID-19 severity. Five severity groups: healthy (0), mild (1–3), moderate (4–5), severe (6–7), and fatal (8).

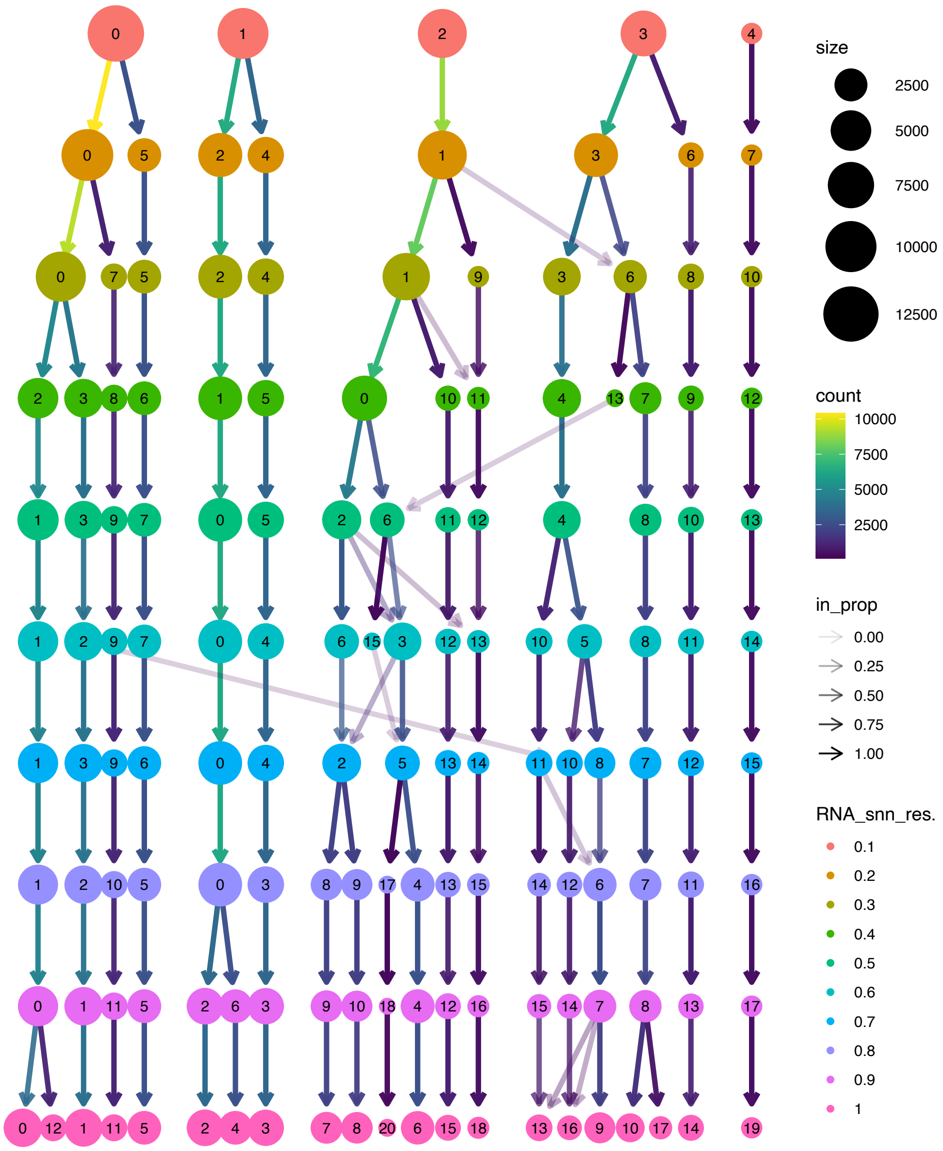


**Supplementary Figure 2. Visualization of cell movements in terms of multiple resolutions.** Clustree was used to visualize and define optimal resolution for cell clustering. For clustering, the resolution parameter varied from 0.1 to 1 with an increasing of 0.1 for every step.

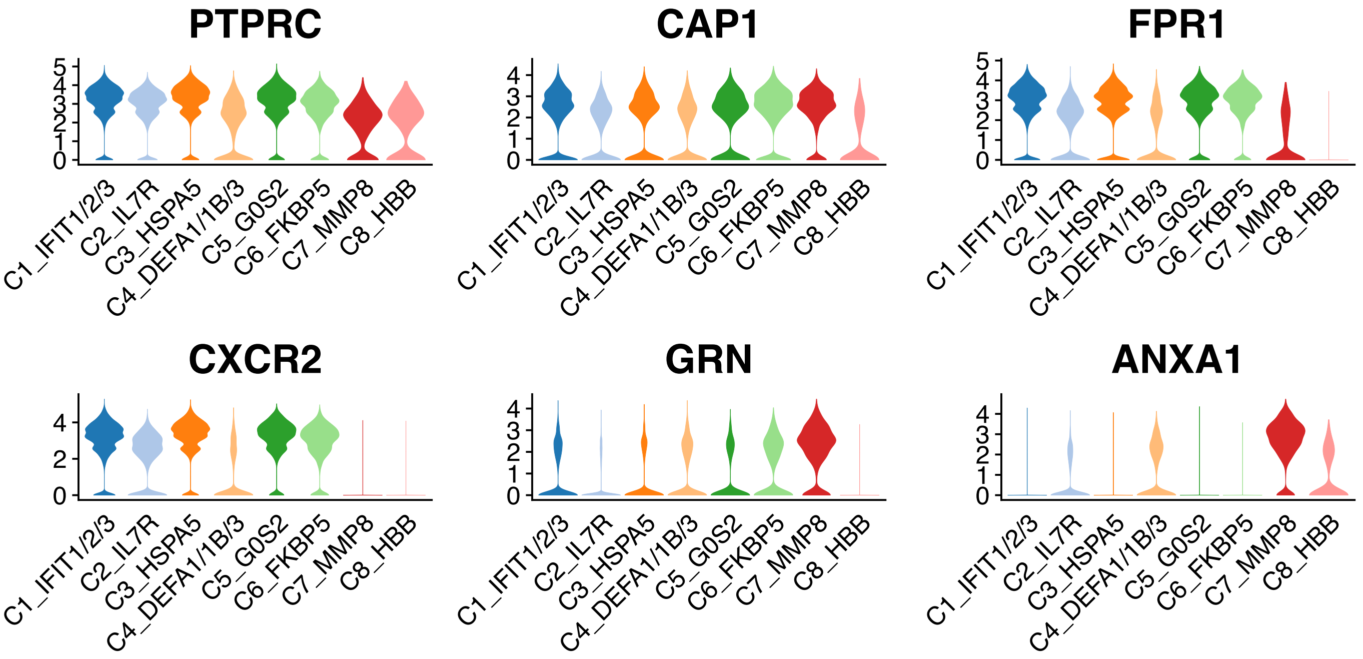


**Supplementary Figure 3. Expressions of selected ligands and receptors.**
Violin plots represent expressions of selected ligands and receptors in Figure 5. Significant gene expression analysis was performed by comparing a relative subtype to all the rest of subtypes. * p value < 0.05; ** p value < 0.01; *** p value < 0.001; N.S.: no significant.

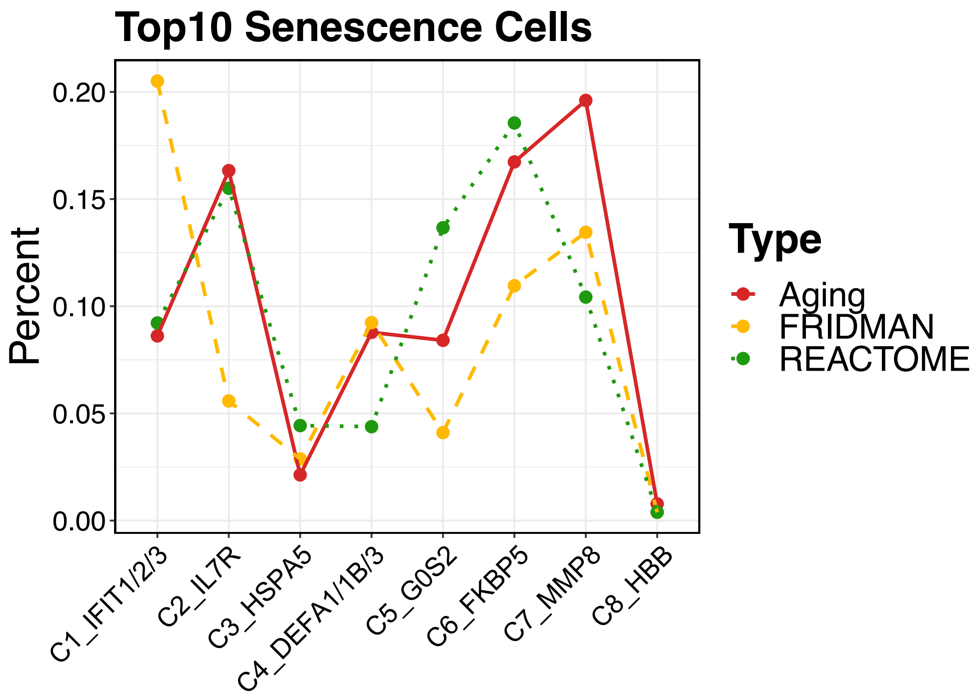


**Supplementary Figure 4. Cell senescence status across eight neutrophil subtypes.** The percentages of each cluster were calculated based on the senescence cells from a cluster. While the senescence cells were defined base average expression of the three genes sets (i.e., aging, fridman, and reactome). In detail, top 10% cells of a given cluster were defined as senescence cells based the average expression levels.

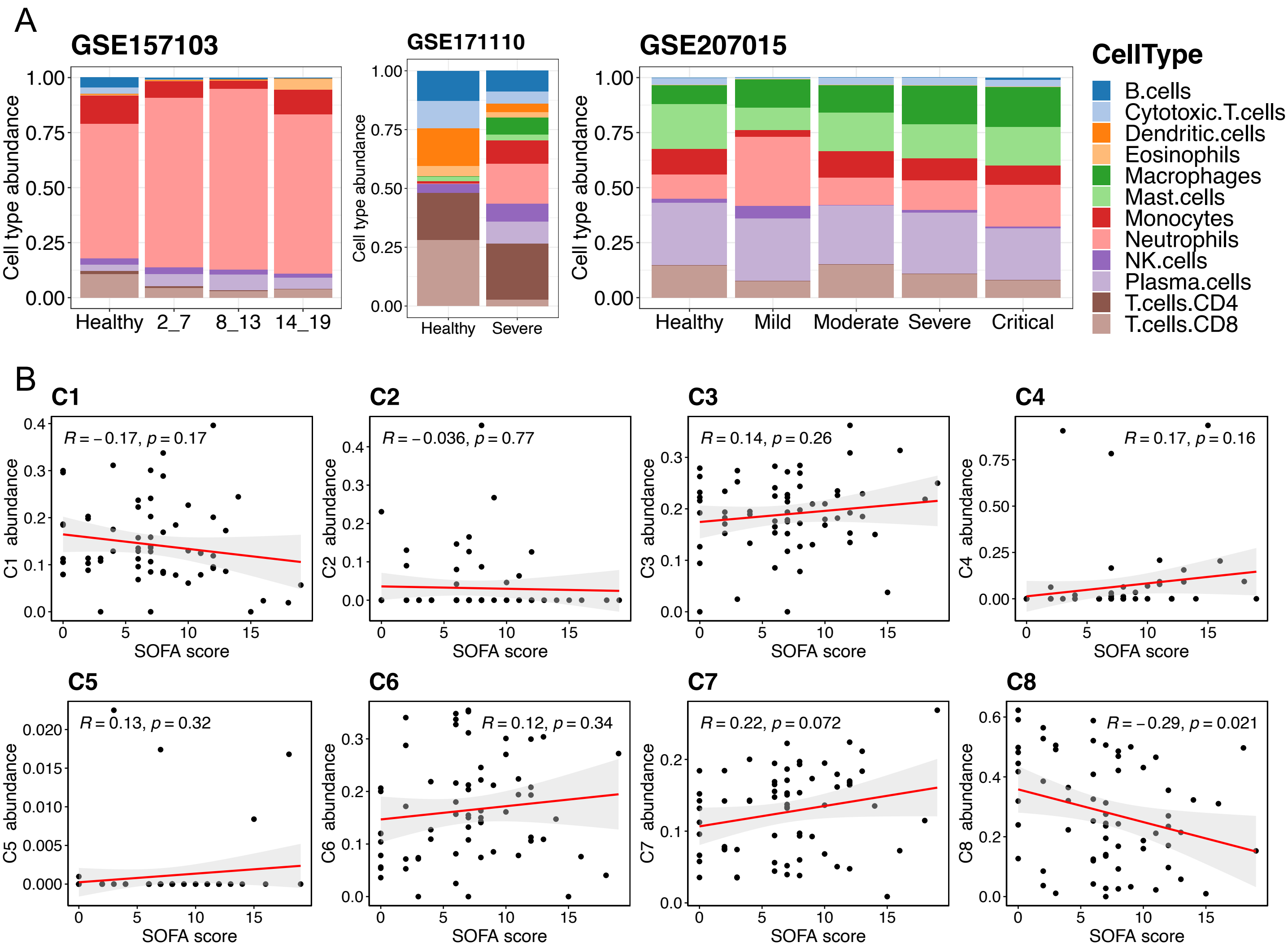


**Supplementary Figure 5. Immune cell type abundance and correlation between neutrophil subtypes and SOFA score.**
(A) Cell type abundance estimation for GSE157103, GSE171110, and GSE207015 datasets using modified LM22 reference. (B) Pearson correlation between the eight neutrophil subtype fractions and SOFA scores using GSE157103 dataset. SOFA: sequential organ failure assessment.

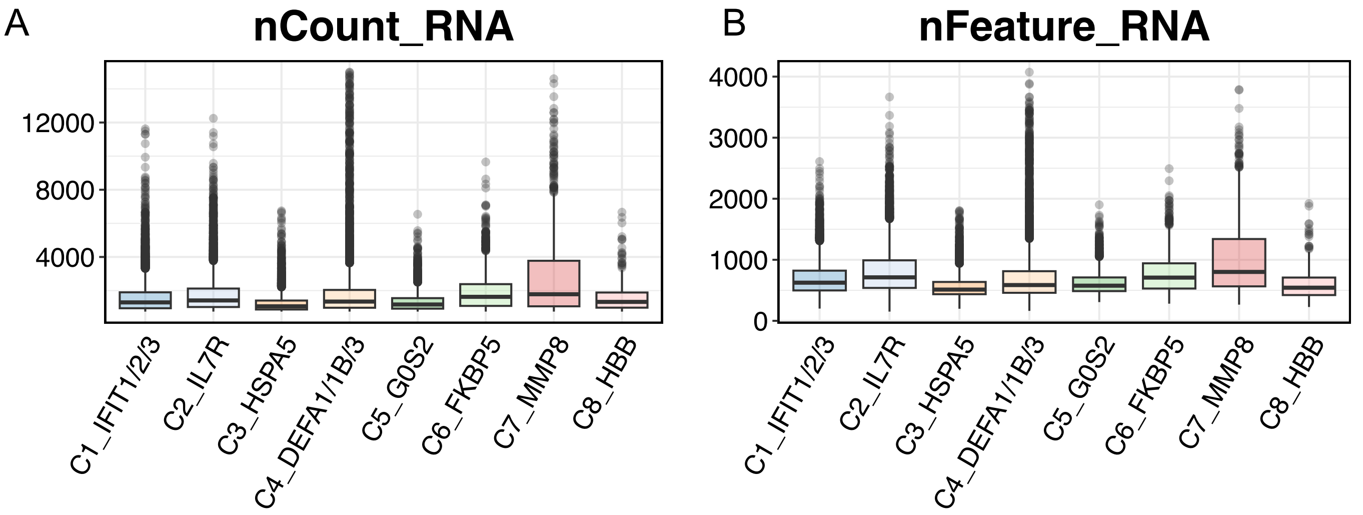


**Supplementary Figure 6. Statistics of the eight characterized subtypes of neutrophils.** For each cell, box plots indicate total numbers of (A) UMI counts and (B) gene across the defined eight subsets of neutrophils, respectively.
